# Nighttime and daytime scenarios in Darwinian liposome evolution under UV-driven natural selection

**DOI:** 10.1101/2025.08.25.671844

**Authors:** Ben Turner, Brian Davies, Gennady Fiksel, Vladimir Subbotin

**Author notes:** Address correspondence to: Vladimir Subbotin, Arrowhead Pharmaceuticals, Madison, WI 53719, USA.

## Abstract

Previously, we presented a hypothesis on Darwinian evolution of liposomes that relies solely on natural and ever-present phenomena: the day-night cycle of solar UV radiation, gravity, and the release of amphiphiles in aqueous media. We demonstrated the protective ability of certain ferric salts typical for Archean waters, notably iron trichloride and ferric ammonium citrate, against liposome destruction by short-wavelength UV by shielding the liposomes with cuvettes filled with ferric salts. In the present study, we investigate the interaction with UV for the liposomes with negative buoyancy that sink naturally in ferric salt solutions. We examined two scenarios: 1) liposomes submerged in the FeCl_3_ solution prior to the UV exposure, a scenario mimicking the nighttime submergence; and 2) submergence under UV exposure, a relevant daytime scenario. We use custom-designed micron-sized, negative-buoyant (heavy), UV-sensitive liposomes. The results showed that for the nighttime scenario, all liposomes were free from UV damage when given sufficient time to submerge in a protective ferric salt solution. Given that larger and heavier liposomes submerge faster, we hypothesize that simple sugar molecules, such as ribose, which is a vital component of RNA, can serve as the proposed heavy molecules in Origin of Life scenarios. Conversely, the daytime scenario results indicate that liposomes slowly submerging under UV exposure sustain some damage. The results of these experiments, which aim to simulate natural events, showed that negative buoyancy and short-wavelength UV radiation could serve as natural drivers in the novel scenario of Darwinian liposome evolution.

## Introduction

The recognized scenarios on the Origin of Life on Earth, e.g., RNA First [1], Hydrothermal Vents [2], Metabolism First [3], assume the occurrence of multiple events that must align strictly consecutively in time and space, i.e., appear as a multistage process. One example is the RNA World hypothesis, i.e., the synthesis of purine and pyrimidine sugars, followed by the synthesis of nucleosides with subsequent polymerization, recombination, folding, and finally, the emergence of the first RNA replicase [4]. However, such co-occurrence of multistage processes raises strong skepticism. For example, A.I. Oparin [5] argued that “the probability that system A will necessarily change to produce system B is very low. But since the probabilities are to be multiplied rather than summed, any multistage process (even the appearance of complex organic substrates) seems to be extremely improbable.” Similarly, L.E. Orgel [6] expressed a strong skepticism regarding spontaneous co-occurrence of multistage processes: “In my opinion, there is no basis in known chemistry for the belief that long sequences of reactions can organize spontaneously, and every reason to believe that they cannot.”.

Additionally, ribozymes and/or RNA cannot survive and evolve by themselves without encapsulation by liposomes, but simultaneous co-emergence in time and space of two events (the ribozymes and the liposomes) appears implausible. In our opinion, one of them must emerge first, survive, and propagate for a long time in abundance. Ribozymes cannot survive and evolve by themselves without liposomes. A logical question is: can liposomes emerge, survive, and then evolve/propagate on their own?

Previously, we presented a hypothesis on Darwinian evolution of liposomes that relies solely on natural and ever-present phenomena: the day-night cycle of solar UV radiation, gravity, and the unique ability of amphiphiles to self-assemble into liposomes, a phenomenon which is the foundation of the Lipid First hypothesis [7-10].

Our hypothesis [11] considers the Archean water-air interface to be a spatial plane, crucial for abiogenesis, in which the ascending amphiphiles form Langmuir layers, bilayers, and liposomes. At the same time, the integrity of liposomes can be strongly affected by solar UV, resulting in their destruction [12-15]. Following the thoughts of Martin Rutten: “One of the many paradoxes encountered in the early history of life lies in the fact that the same rays of the Sun which formed the building blocks of the molecules of life were lethal for life. Early life had, therefore, only limited environmental possibilities. It could be survived only when shielded by a thick layer of water…”[16], we hypothesized that some liposomes encapsulate a solute heavier than the surrounding media and, by descending from the water-air interface, are protected from UV because of the UV absorption in water. We also calculated the depth-vs-time dynamics of the submergence, which depends on known physical characteristics: liposomal size, specific gravity, and viscosity of the media [11]. However, the UV-shielding ability of the presumable Archean water strongly depends on its composition and remains a subject of experimental tests.

It is known, and we demonstrated that pure water has a very weak UV absorption and must contain certain impurities to exhibit shielding ability. Therefore, we investigated [17] the attenuating ability of several ferric salts typical for Archean waters. We showed that iron trichloride and ferric ammonium citrate aqueous solutions, at a concentration of 2.5 g/L, attenuate the UV intensity by a factor of 1000 at a submersion depth of 6.3 mm and 7.4 mm, respectively.

In the follow-up experiments [18], we examined 1) what part of the UV spectrum produces a damaging effect on UV-sensitive liposomes and investigated 2) whether solutions of iron trichloride and ferric ammonium citrate can reduce or eliminate the UV-damaging effect and protect liposomes. In that study, we did not directly submerge liposomes into the solutions but instead shielded them with UV-transparent cuvettes filled with ferric salt solutions. We used specially designed UV-sensitive liposomes (Encapsula NanoSciences, Brentwood, USA). The results of this study demonstrated that 1) liposomes can be destroyed by short UV wavelengths (</ = 254 nm) after the UV exposure as short as 6 min, 2) long-wave UV radiation (∼315–400 nm), including atmosphere-filtered Sun UV, do not destroy liposomes, and 3) a thin layer (10 mm) of ferric salt solutions, iron trichloride and ferric ammonium citrate, completely protect the liposomes from short-wavelength UV. These results reinforce our hypothesis [11] that the Sun’s UV radiation and gravity could be the major selection forces for abiogenesis.

### Present study – objectives and brief summary

The goal of the present study is to examine two scenarios: 1) liposomes submerged in the FeCl_3_ solution prior to the UV exposure, a scenario mimicking the nighttime submergence; and 2) liposomes submerged in the FeCl_3_ solution under UV exposure, a relevant daytime scenario. We use custom-designed micron-sized, negative-buoyant (heavy), UV-sensitive liposomes. The results showed that for the nighttime scenario, all liposomes were free from UV damage when given sufficient time to submerge in a protective ferric salt solution. Given that larger and heavier liposomes submerge faster, we hypothesize that simple sugars, such as ribose, which is a vital component of RNA, can serve as the proposed heavy molecules in Origin of Life scenarios. Conversely, the liposomes submerged in pure water were completely destroyed. On the other hand, the daytime scenario results (liposomes submerged in the FeCl_3_ solution under UV exposure) indicate that liposomes slowly submerging under UV exposure sustain some damage, although not as strong as in pure water.

The results of these experiments, which aim to simulate natural events, showed that negative buoyancy and short-wavelength UV radiation could serve as natural drivers in the novel scenario of Darwinian liposome evolution.

## Material and Methods

### Liposomes

We used large polymerizable UV-sensitive multilamellar micron-sized liposomes composed of Dibehenoyl-sn-glycero-3-phosphocholine (DBPC) and a 10,12-Pentacosadiynoic acid (PCDA) with a DBPC: PCDA molar ratio of 80:20, at a total lipid concentration of 7 mM (Encapsula NanoSciences, Brentwood, TN, USA). The mechanisms of liposomal UV sensitivity and UV destruction were described in the previous publication [18]. Briefly, the PCDA is a polymerizable unsaturated fatty acid. PCDA lipids adjacent to each other polymerize upon UV radiation, changing their color from the original white to blue [19]. For the negative buoyancy experiments, liposomes were manufactured in water with sucrose, allowing them to reach a specific gravity of 1.014 g/cm^3^. According to the manufacturer’s description, liposomes were “micron-sized”.

### UV source and measurement of UV intensity

We used a mercury low-pressure 15-Watt Spectroline Germicidal UV Sterilizing lamp (General Tools, USA), with an intensity in the UVC range (245 nm) of 9,500 μW/cm^2^ at a distance of 30 mm from the lamp (UV512C Light Meters, General Tools, USA).

### UV shielding medium

Based on our previous observations, we used a water solution of iron trichloride at a concentration of 2.5 g/L, with a specific gravity of 1.008 g/cm^3^.

### Assessment of liposomal damage

The liposomal damage was assessed visually and imaged with a KEYANCE BZ-X810 microscope at a x400 magnification. It was also quantified by measuring the absorbance of the scanning light with a wavelength of 645 nm emitted by a BioTek Synergy Neo2 Hybrid Multimode Reader (Agilent, Santa Clara, CA, USA). The absorbance data were acquired by a Perkin Elmer Synergy Neo2 plate reader, as previously described [18].

### Experimental setup for measurement of liposomes’ size and the rate of submersion

The liposome size assessment was done using the inverted laser scanning confocal microscope LSM 980, utilizing the scanning light at 488 nm. The liposome size was quantified with the ZEN program measuring tool utilizing 15 µm calibration beads as a reference point. For the submersion rate measurement, the scanning images were acquired every 30 minutes for 18 hours.

The experimental setup consisted of a 96-well glass-bottom plate (Cellvis, USA), two wells of which were prefilled with 270 μL of water containing 15-micron calibration beads. The plate was positioned on a stage of the inverted laser scanning confocal microscope (LSM 980). As soon as beads were identified at the bottom of the wells, 30 μL of diluted liposomes (1:10 in water) were slowly layered on top of the central point of the solutions, using previously mounted micro tubing, to avoid mechanical disturbance of the solutions. An alternating confocal scanning of the bottom planes of two wells was performed every 30 minutes for 18 hours.

To study the effects of UV radiation on liposomes, we used a plastic-bottom 96-well plate (Greiner, Austria). In all experiments, 270 μL of solutions were added to the wells (iron trichloride or water solutions), making the solution depth 10 mm. To examine the nighttime scenario, 30 μL of undiluted liposomes were layered on top of 270 μL of iron trichloride solution or water 14 hours prior to UV radiation. To examine the daytime scenario, 30 μL of undiluted liposomes were slowly (5 μL/min, 30 μL in 6 minutes) layered on top of 270 μL of iron trichloride solutions during 6 minutes of UV radiation.

To prevent uneven distribution of liposomes and mechanical disturbance, as well as to avoid UV light obstruction, we employed a specialized design for the slow delivery of liposomes during UV radiation. The design comprises three 5 cm 33 G hypodermic tubings, extended to the center of targeted wells, with tips curved so that they touch the surface of the solution (Figure 1).

**Figure 1.**
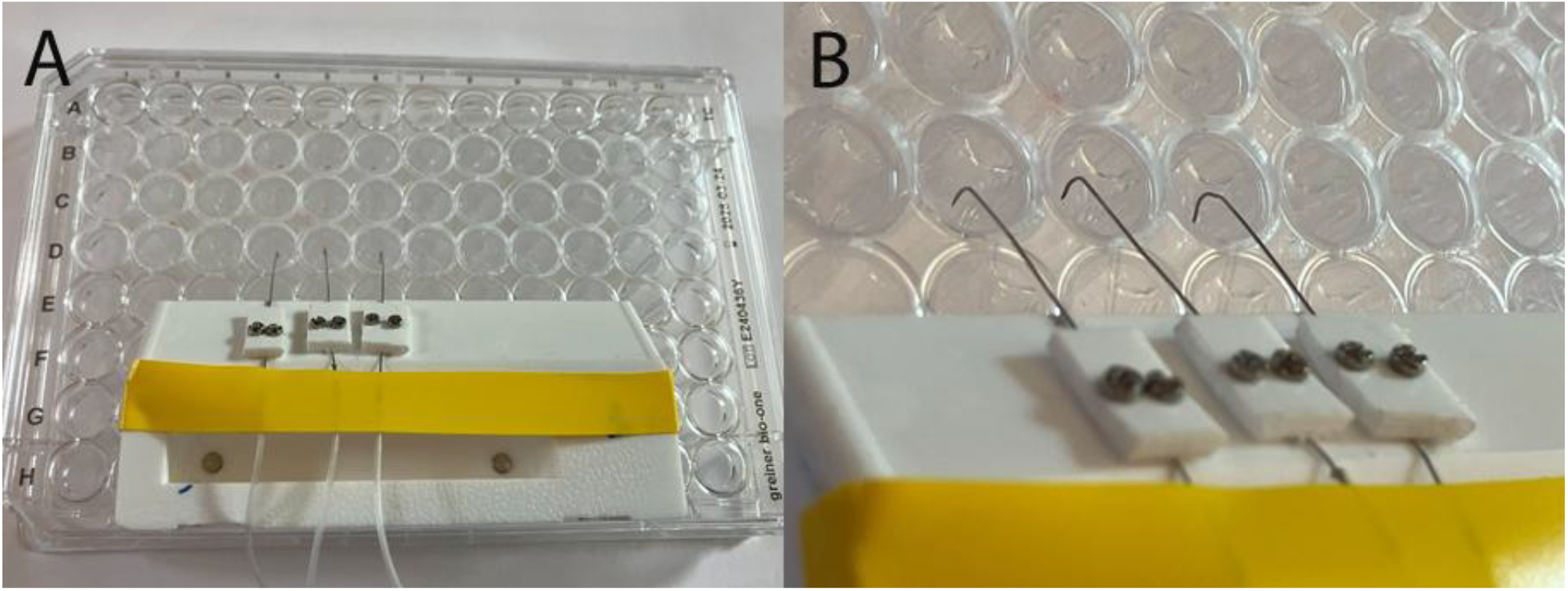
A devise for slow delivery of liposomes during UV radiation (daytime scenario). **A** – a photo of the entire plate with the device, **B** – a magnified portion showing the curved tips.

We used three 50 μL Hamilton syringes and a Harvard Apparatus syringe pump for slow delivery of liposomes during UV radiation. This design provides a smooth entrance of the liposome into the solution, thus removing the effect of the initial velocity on the submergence time, which then remains governed solely by the gravity, buoyancy, and the solution viscosity.

The UV lamp was positioned 30 mm above the plates, making the UVC intensity at the targets 9,500 μW/cm^2^. The 6 minutes UV radiation time was shown in previous experiments to be sufficient for complete liposome destruction [18]. Liposomal damage was assessed visually and quantified by measuring the absorbance of the scanning 645 nm light emitted by a BioTek Synergy Neo2 Hybrid Multimode Reader (Agilent, Santa Clara, CA, USA). The absorbance data was acquired by a Perkin Elmer Synergy Neo2 plate reader as previously described [18].

## Results

### Liposomes’ size distribution and the submersion time

An example of liposomal size measurement is shown in Figure 2. Analysis of scanning images taken every 30 minutes for 18 hours showed that all liposomes had settled on the bottom of the wells in 11.5 hours.

**Figure 2.**
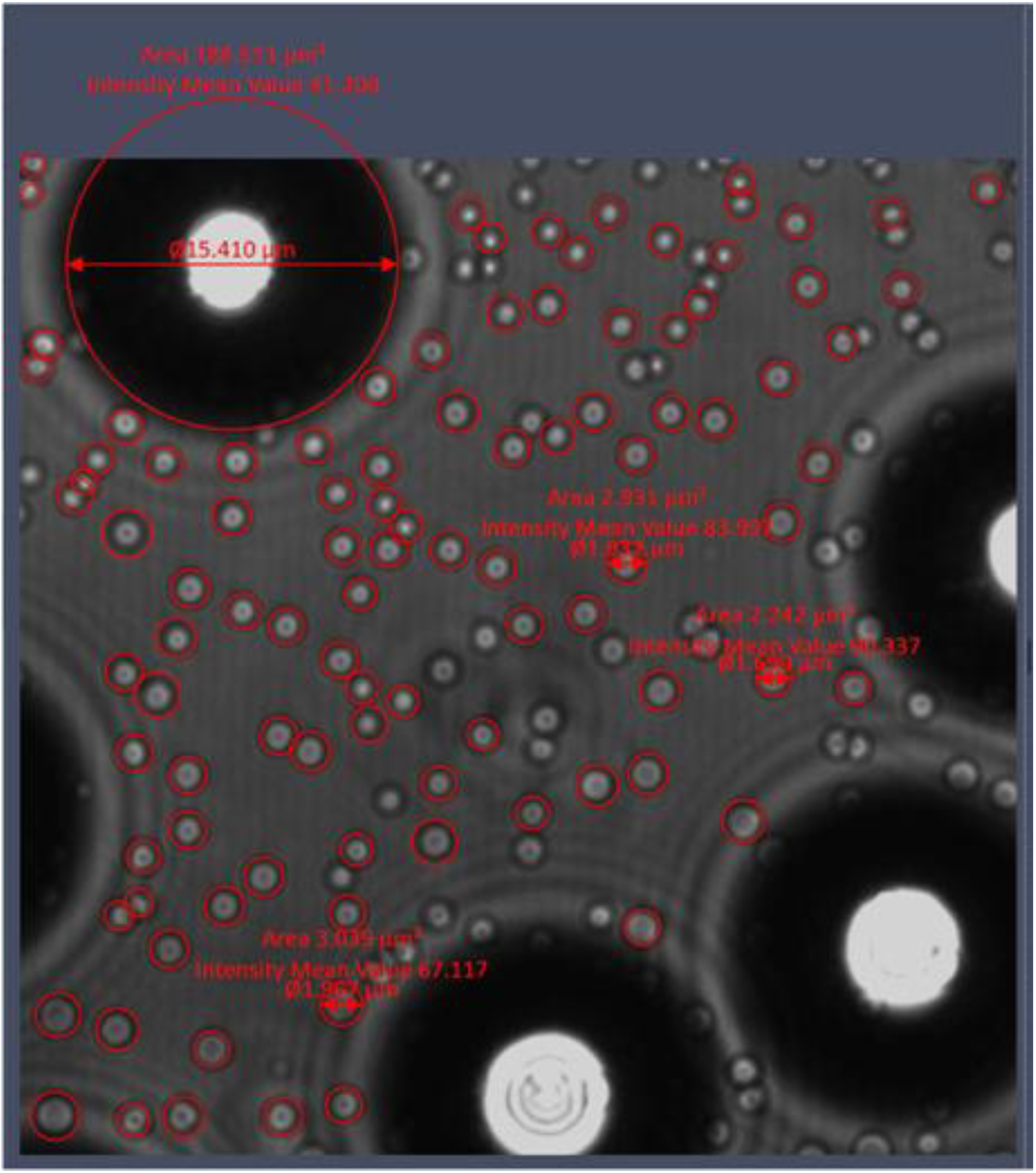
Liposomes’ image with the confocal microscope (LSM 980 utilizing the scanning light at 488 nm). A hundred liposomes were measured, using 15-micron calibration beads as a reference point.

The size distribution of the liposomes is shown in Figure 3. The liposome average diameter is 1.64 μm, and the size distribution standard deviation is 0.23 μm.

**Figure 3.**
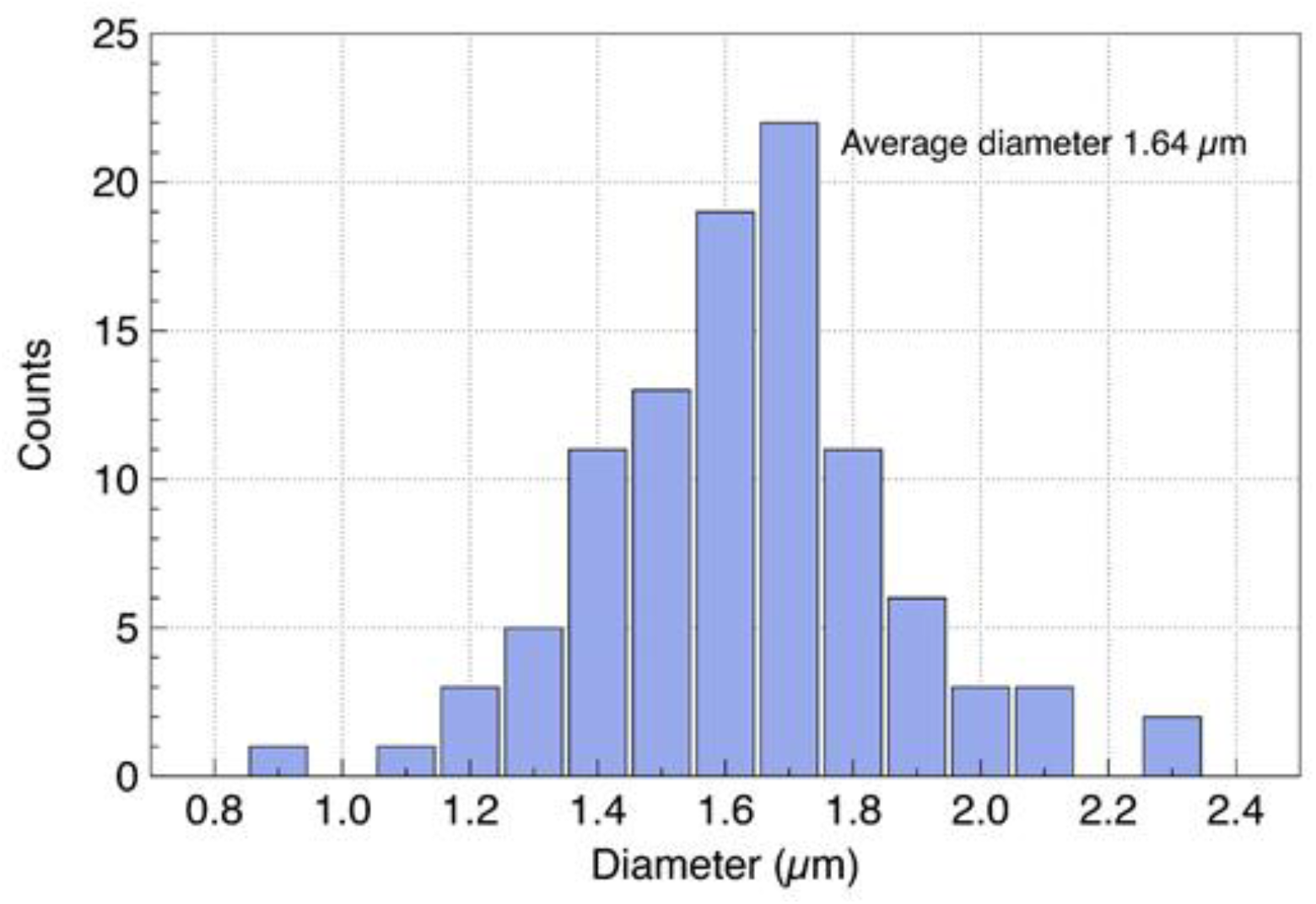
Liposomal size distribution.

### Investigation of the nighttime and daytime scenarios of liposome submergence

A qualitative visualization of the UV damage is illustrated in Figure 4. As mentioned before, the damage is associated with the intensity of the blue color of the liposomes after the UV exposure. The results showed that liposomes added 14 hours prior to UV radiation to the wells filled with pure water were destroyed entirely (wells D 10-12) as indicated by the intense blue tint, while liposomes added 14 hours prior to UV radiation to the wells with iron trichloride solution were completely protected (wells D 7-9). Liposomes that were added to the wells with iron trichloride solution during UV radiation also underwent damage, as indicated by a color change, although it was less than that for liposomes shielded by pure water (Figure 4).

**Figure 4.**
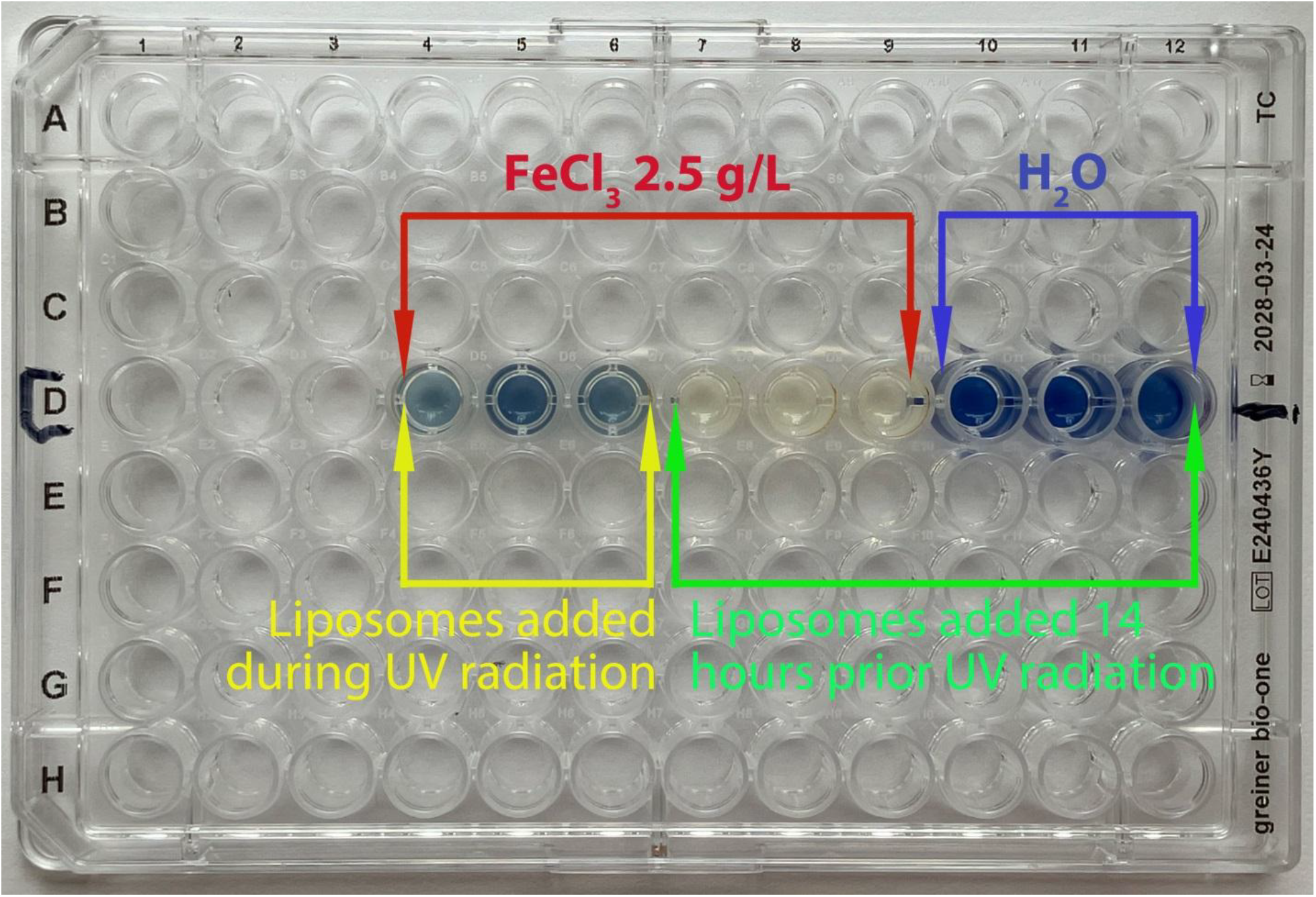
A photo of the experimental design with a 96-well plate after 6 minutes of UV radiation. Liposomes “shielded” by water are completely destroyed (D 10, 11, 12), while liposomes shielded by FeCl_3_ solution appeared to be completely protected (D 7, 8, 9). Liposomes added to FeCl_3_ solution during UV radiation also appear to be destroyed (D 4, 5, 6), although the color shift is visibly less than that for the unshielded liposomes (H_2_O wells).

This visual evaluation was aided by measuring the absorbance of the scanning 645 nm light emitted by a BioTek Synergy Neo2 Hybrid Multimode Reader, as previously described [18] (Figure 5).

**Figure 5.**
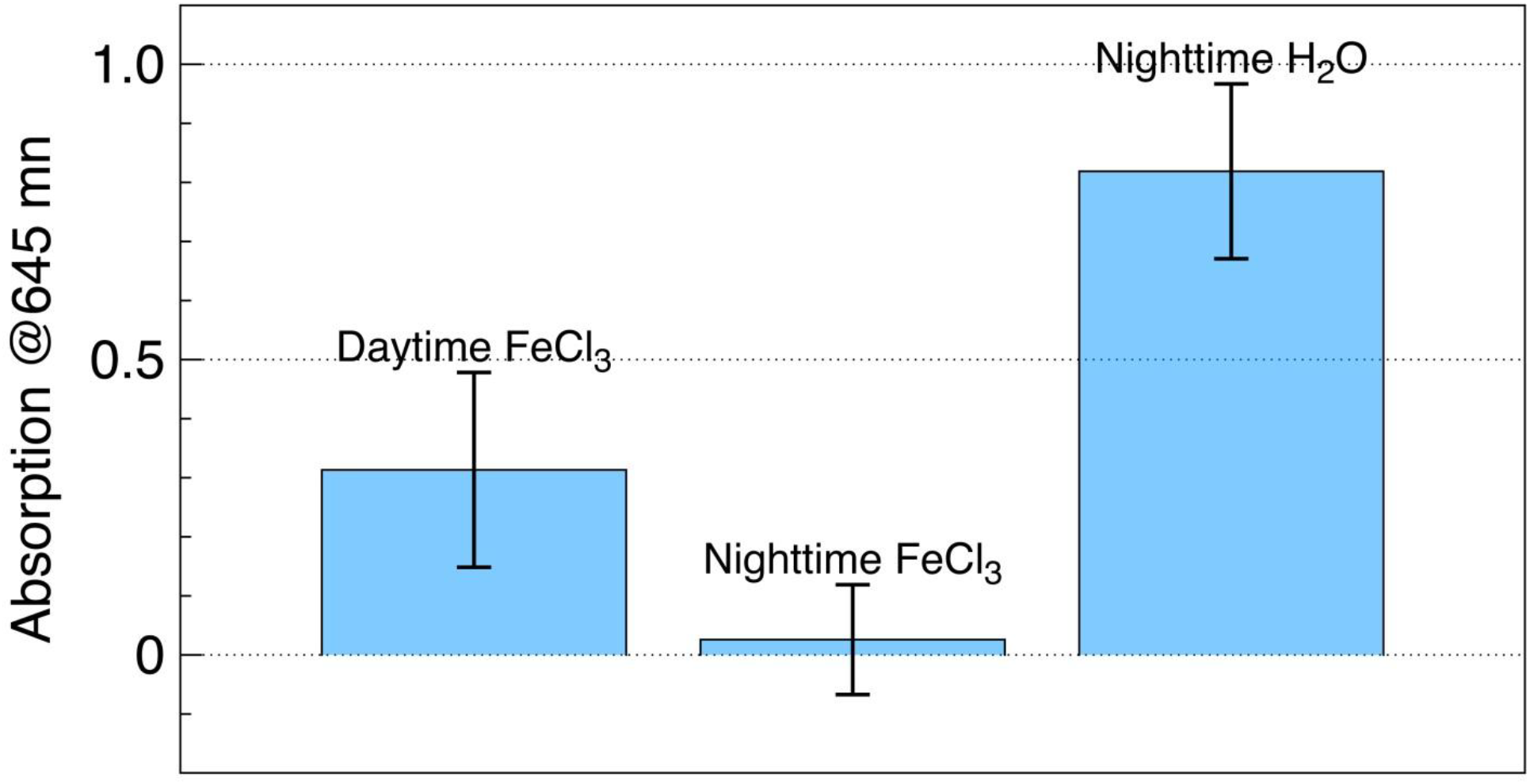
Color shift measured via absorption of 645 nm light by BioTek Synergy Neo2 Hybrid Multimode Reader.

The UV-induced nighttime absorption change for unprotected liposomes (H_2_O wells) is 0.8, while that for the protected liposomes (FeCl_3_ wells) is nearly zero. The absorption change for the protected liposomes added during UV radiation (FeCl_3_ wells, daytime scenario) is 0.3. That low damage was unexpected, given that at an average submersion rate of ∼1 mm/hr, the liposomes could only sink about 0.1 mm in 6 minutes, which is not sufficient to be protected from UV. One possible explanation is that larger liposomes, originating either from the distribution tail shown in Figure 3, or from the fusion or clamping of smaller liposomes, can submerge faster and may gain at least partial protection, which could explain the lower damage observed.

## Summary and conclusion

The results of these experiments, aiming to simulate natural events, showed that negative buoyancy and UV radiation are sufficient as natural drivers in a novel scenario of Darwinian evolution of liposomes. It was established that liposomes added prior to the UV radiation (nighttime scenario) and completely submerged into ferric salt aqueous solution (FeCl_3_, 2.5 g/L) are totally protected from the UV damage, while liposomes submerged into pure H_2_O are destroyed. Liposomes added during UV exposure (daytime scenario) are destroyed even in ferric solution because of the insufficient rate of submersion. Note here that to obtain measurable data in a limited amount of time, a high intensity of UV light was used. In particular, the UVC intensity was about 100 W/m^2^, which is about 2 orders of magnitude higher than that currently observed in nature. Thus, the daytime formation and subsequent submersion and protection of liposomes under natural conditions cannot be dismissed. Further experiments on nighttime/daytime scenarios of liposomes’ survival are in progress.

## Notes

### Competing Interest Statement

The authors have declared no competing interest.

## References

1 Pressman, A. et al. (2015) The RNA world as a model system to study the origin of life. Current biology 25 (19), R953–R963

2 Martin, W. et al. (2008) Hydrothermal vents and the origin of life. Nature Reviews Microbiology 6 (11), 805–814

3 Wächtershäuser, G. (2006) From volcanic origins of chemoautotrophic life to Bacteria, Archaea and Eukarya. Philosophical Transactions of the Royal Society B: Biological Sciences 361 (1474), 1787–1808

4 Le Vay, K. and Mutschler, H. (2019) The difficult case of an RNA-only origin of life. Emerging Topics in Life Sciences 3 (5), 469–475

5 Oparin, A. (1974) A hypothetic scheme for evolution of probionts. Origins of life 5 (1), 223–226

6 Orgel, L.E. (1998) The origin of life—a review of facts and speculations. Trends in biochemical sciences 23 (12), 491–495

7 Morowitz, H.J. et al. (1988) The chemical logic of a minimum protocell. Origins of Life and Evolution of the Biosphere 18 (3), 281–287

8 Luisi, P.L. et al. (1999) Lipid vesicles as possible intermediates in the origin of life. Current opinion in colloid & interface science 4 (1), 33–39

9 Stano, P. and Luisi, P.L. (2010) Achievements and open questions in the self-reproduction of vesicles and synthetic minimal cells. Chemical Communications 46 (21), 3639–3653

10 Segré, D. et al. (2001) The lipid world. Origins of Life and Evolution of the Biosphere 31 (1), 119–145

11 Subbotin, V. and Fiksel, G. (2023) Exploring the Lipid World hypothesis: A novel scenario of self-sustained Darwinian evolution of the liposomes. Astrobiology 23 (3), 344–357

12 Putvinskij, A. et al. (1977) Study of the effect of ultraviolet light on biomembranes. 6. Studia biophysica 64 (1), 17–32

13 Potapenko, A.Y. et al. (1972) Investigation into the UV-radiation effect on biological membranes. The change in electric conductivity of bimolecular phospholipid membranes. Doklady Akademii Nauk SSSR, Seriya Fizicheskoj Khimii 202 (4), 882–885

14 Vladimirov, Y.A. et al. (1986) Electrical stability of artificial membranes. Gen. Physiol. Biophys 5, 231–242

15 Putvinsky, A. et al. (1979) Electric breakdown of bilayer phospholipid membranes under ultraviolet irradiation-induced lipid peroxidation. FEBS letters 106 (1), 53–55

16 Rutten, M.G. (1971) Origin of life by natural causes, American Elsevier

17 Subbotin, V. and Fiksel, G. (2023) Aquatic Ferrous Solutions of Prebiotic Mineral Salts as Strong UV Protectants and Possible Loci of Life Origin. Astrobiology 23, 741–745

18 Turner, B. et al. (2025) Protection of liposomes by ferric salts against the UV damage and its implications for the origin of life. Frontiers in Astronomy and Space Sciences 12, 1566396

19 Ahn, D.J. et al. (2009) Rational design of conjugated polymer supramolecules with tunable colorimetric responses. Advanced Functional Materials 19 (10), 1483–1496

